# Pushing the boundaries of MEG based on optically pumped magnetometers towards early human life

**DOI:** 10.1101/2023.10.28.564455

**Authors:** Pierre Corvilain, Vincent Wens, Mathieu Bourguignon, Chiara Capparini, Lauréline Fourdin, Maxime Ferez, Odile Feys, Xavier De Tiège, Julie Bertels

## Abstract

Characterizing the early development of the human brain is critical from both fundamental and clinical perspectives. However, existing neuroimaging techniques are either not well suited to infants or have limited spatial or temporal resolution. The advent of optically pumped magnetometers (OPMs) has revolutionized magnetoencephalography (MEG) by enabling wearable and thus more naturalistic recordings while maintaining excellent sensitivity and spatiotemporal resolution. Nevertheless, its adaptation to studying neural activity in infancy poses several challenges. In this work, we present an original close-to-scalp OPM-MEG setup that successfully recorded brain responses to sounds in newborns. We exposed one-month-old infants to continuous streams of tones and observed significant evoked responses, which peaked ∼250 ms poststimulus at bilateral auditory cortices. When tones were presented at a steady fixed pace with an oddball tone every fourth tone, significant neural responses were found both at the frequency of the standard tones (3 Hz) and of the oddball tones (0.75 Hz). The latter reflects the ability of the newborn brain to detect auditory change and synchronize to regular auditory patterns. Additional analyses support the added value of triaxial OPMs to increase the number of channels on small heads. Finally, OPM-MEG responses were validated with those obtained from the same participants using an adult-sized cryogenic MEG. This study demonstrates the applicability of OPM-MEG to study early postnatal periods; a crucial step towards future OPM investigations of typical and pathological early brain development.

## Introduction

During the early stages of life, the human brain undergoes major changes, with the emergence of perceptual, motor, language, social and cognitive skills. Characterizing the neurophysiological processes underlying the development of these early abilities is crucial to understanding brain maturation and brain pathologies.

However, existing brain imaging techniques are limited in investigating early brain development. While wearable systems such as electroencephalography (EEG) and near-infrared spectroscopy (NIRS) can be adapted to small heads and enable naturalistic recordings, they have limited spatial (EEG) or temporal (NIRS) resolution. Moreover, fixed systems such as functional magnetic resonance imaging (MRI) and cryogenic magnetoencephalography (MEG) have good spatial (both) and temporal (MEG) resolutions but are not adapted to—and often not well tolerated by—infants. Thus, in practice, infants are often restrained or sedated when using these techniques (Copeland et al., 2021).

The excellent temporal and very good spatial resolution of MEG (Baillet, 2017) can be leveraged by having the sleeping infant rest their head on an MEG array (either using adult-size helmets in the supine position (Huotilainen et al., 2003) or concave arrays designed to fit maternal wombs (Draganova et al., 2007; Holst et al., 2005; Lengle et al., 2001; Muenssinger et al., 2013)). However, only partial head coverage can be obtained with this approach unless several recordings are performed (Y.-H. Chen et al., 2019). Some efforts have been made to perform whole-head MEG scans with infants. When adult-sized cryogenic MEG systems are combined with a foam halo, the infant’s head can be supported and centered in the helmet (Clarke, Bosseler, et al., 2022; Clarke, Larson, et al., 2022), but the resulting large brain-to-sensor distance is detrimental to both the signal-to-noise ratio (SNR) and spatial resolution (Wens, 2023). To address this issue, infant-sized cryogenic MEG systems have been developed (Y.-H. Chen et al., 2019). Still, these systems are extremely expensive and not usable in other age ranges, so very few systems are available worldwide (Feys & De Tiège, 2024).

Optically pumped magnetometers (OPMs) are novel magnetic field sensors that offer considerable flexibility while preserving the outstanding sensitivity of cryogenic magnetometers to the subtle magnetic fields produced by neural currents (Boto et al., 2017). In contrast to cryogenic MEG systems, in which sensors are fixed in rigid helmets, OPMs can be used to form wearable sensors arrays (OPM-MEG) that can be placed on scalp, allowing more natural and flexible scanning situations (Boto et al., 2018; Brookes et al., 2022). In addition, the ensuing reduction in brain-to-sensor distance improves signal amplitude and spatial resolution (Wens, 2023). Crucially, a single OPM-MEG system can be adapted to all head sizes, whereas several dedicated cryogenic MEG systems would be needed. OPM-MEG systems also consume less energy (direct electricity consumption; see Methods section) and are more affordable than cryogenic MEG systems, which is critical for their future dissemination.

The compliance of OPM-MEG has been demonstrated over a wide range of ages, starting from two years old, for both physiological (Hill et al., 2019; Rhodes et al., 2024; Rier et al., 2024) and clinical (Feys, Corvilain, Van Hecke, et al., 2023; Feys et al., 2022, 2024) recordings. However, the use of OPM-MEG in younger subjects poses substantial methodological challenges. First, the surface of alkali OPMs is too warm for prolonged direct contact with the skin because the rubidium inside the sensitive cell needs to be heated to approximately 150°C (Brookes et al., 2022). Therefore, due to lack of hair and skin sensitivity of newborns, efficient heat dissipation is needed to prevent discomfort and injuries. Second, multiple OPMs must be fixed on the small heads of infants to perform whole-head OPM recordings, which requires balancing ergonomics and signal quality. Recently developed triaxial OPMs increase the number of channels without impacting the ergonomics. This enables better denoising (Brookes et al., 2021) and, in infant studies where spatial sampling is limited, measurements in the direction tangential to the scalp may capture neural activity missed by measurements in the radial direction (Boto et al., 2022; Feys, Corvilain, Labyt, et al., 2023).

In this work, we present a wearable OPM-MEG setup that is suitable for recordings starting from birth and we demonstrate its ability to accurately characterize early brain functions in 14 one-month-old healthy neonates. We focused on auditory stimulation, as auditory evoked brain responses have been well described in infants, in particular using EEG (Barnet et al., 1975a; Dehaene-Lambertz, 2000; Kushnerenko et al., 2002; Wunderlich & Cone-Wesson, 2006), and they occur regardless of the newborn’s behavioral state (Martynova et al., 2003) and position with respect to the speaker. We hypothesized that OPM-MEG will be able to accurately detect and localize typical auditory evoked responses as well as neural responses elicited by a fast periodic auditory oddball paradigm. We tested the added value of triaxial OPMs to increase the number of channels on small heads. Finally, we validated the OPM-MEG responses with those obtained from the same participants using an adult-sized cryogenic MEG. Overall, this work provides unprecedented foundation for future OPM-MEG neurodevelopmental studies in very young participants.

## Results

### System design

To balance the requirements of signal quality, adaptability, comfort and heat dissipation, we developed flexible EEG-like caps to which custom 3D-printed OPM holders that maintain the sensors 4 mm above the scalp were sewn (Fig. 1). The caps were equipped with ∼27 small and lightweight alkali OPMs (Gen2/Gen3 QZFM, QuSpin Inc.), which almost entirely covered the scalp. As the newborns were meant to rest on their back, no sensors were placed in the occipital region. The setup was comfortable and light enough to be well tolerated by neonates during the whole experiment while resting on their parent’s lap or chest. This OPM-MEG setup has previously been used by our group with a 5-month-old epileptic infant to study epileptiform discharges <u>(Feys, Corvilain, Bertels, et al., 2023)</u>. Source localizations of the group level responses were enabled by placing the average-sized cap on a 3D printed surface of a template MRI for sensors digitalization and MRI co-registration (see Methods section and Fig. S1 in the Supplemental Material).

**Fig. 1.**
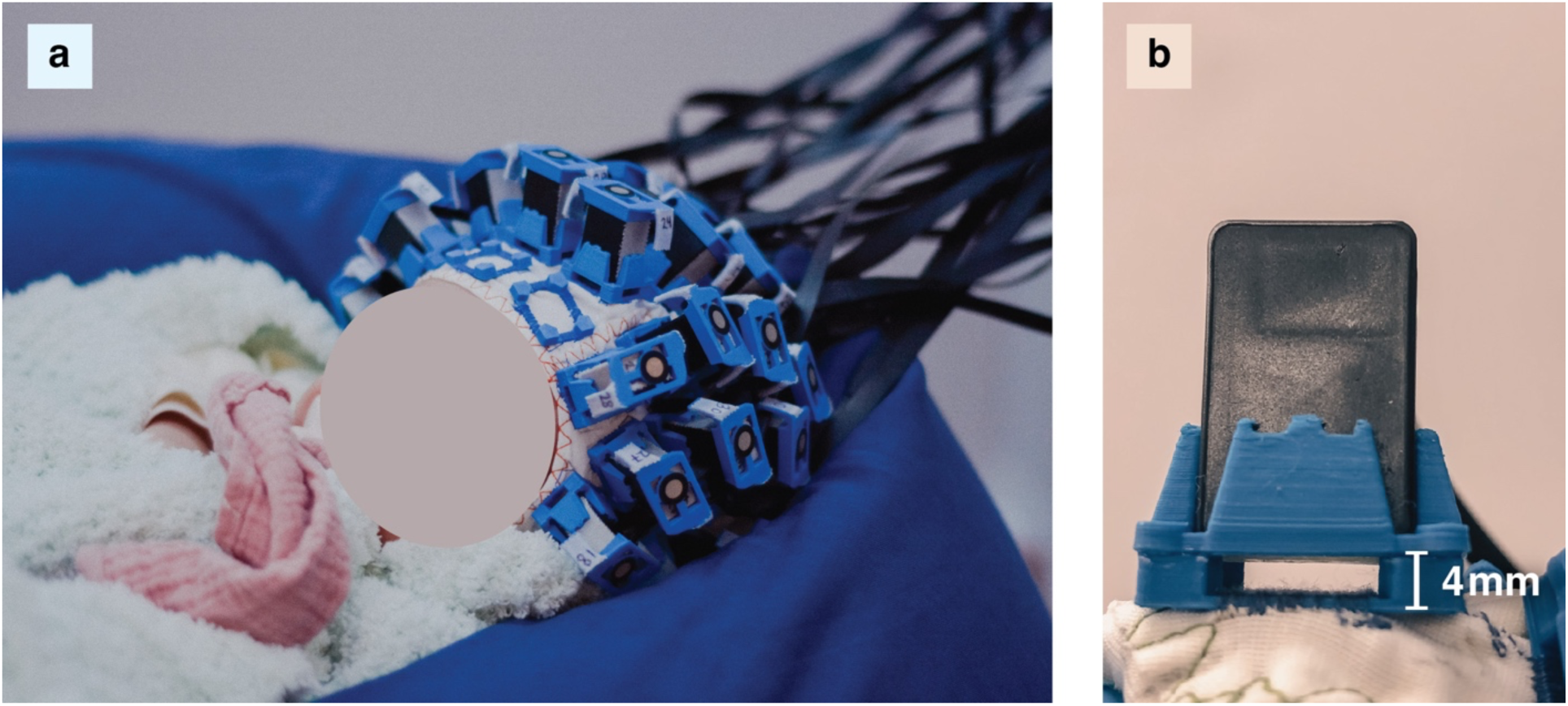
Experimental setup. (**a**) A newborn undergoing the experiment with the OPM-MEG system. (**b**) Close-up view of the sewed-on OPM holders designed on site. The 4-mm spacing prevents direct contact between the warm OPM and the neonate scalps.

### Auditory evoked responses

In a first paradigm, we presented auditory pure tones every 500–600 ms (Fig. 2a) to demonstrate that our OPM-MEG setup can reliably detect evoked responses. The grand-average evoked fields in the radial direction displayed a clear response to the stimuli (Fig. 2b). The associated topographic plot exhibited the expected dipolar pattern above the bilateral temporal regions (Fig. 2c), and the source localization showed activity peaking at the auditory cortices bilaterally (Fig. 2d). This response was statistically significant from 179 ms to 359 ms poststimulus (*p* < 0.0001, maximum statistics) and its root-mean-square peaked at 261 ms. Similar analyses performed at the individual level revealed significant responses in 9 participants.

**Fig. 2.**
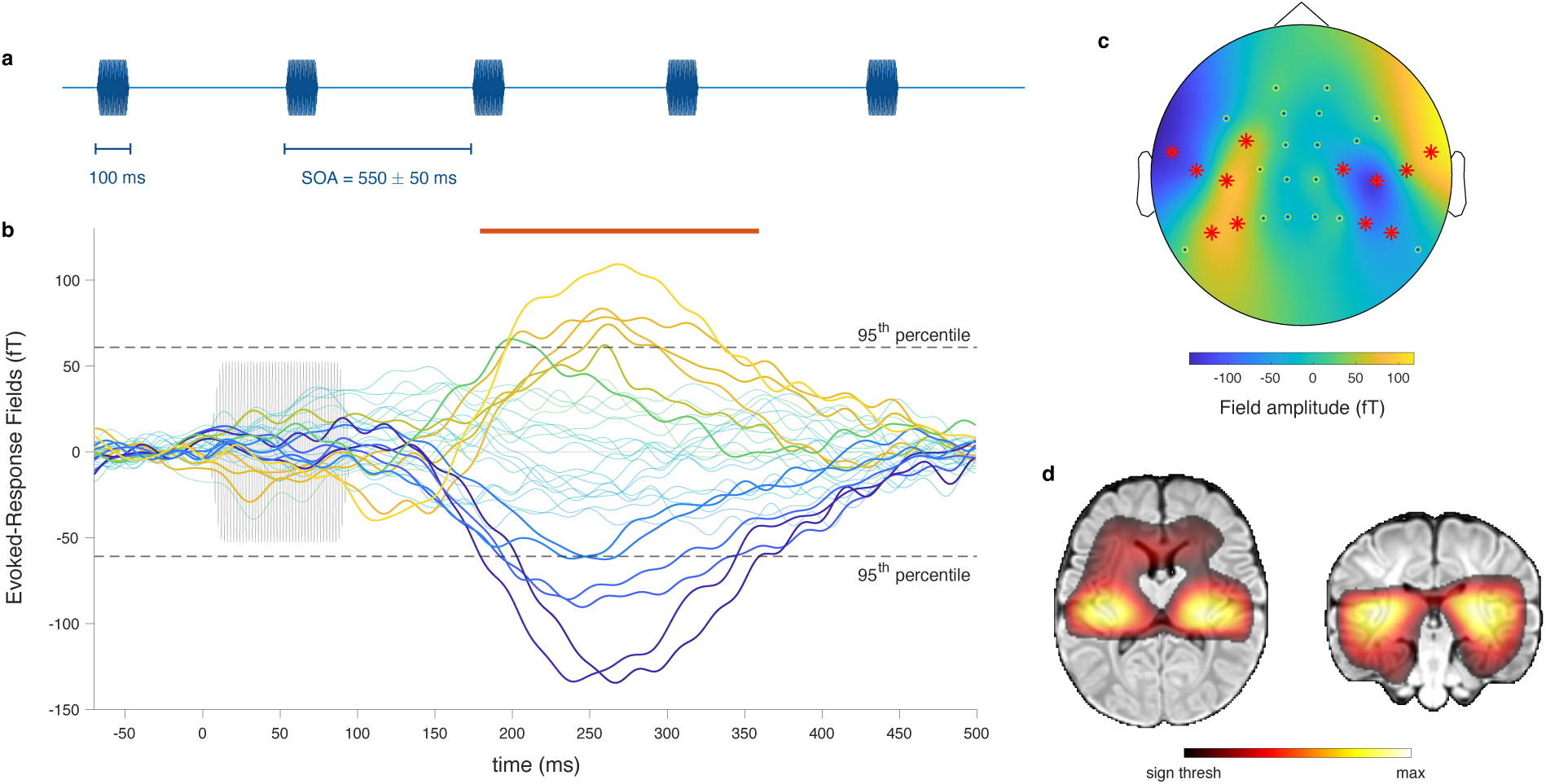
Auditory evoked response paradigm. (**a**) Excerpt from the sound sequence, during which a single pure tone is presented repeatedly with a variable stimulus onset asynchrony (SOA). (**b**) Time course of the group-level evoked magnetic field measured by the OPM radial axes from −70 ms to 500 ms peristimulus. The sound stimulus is displayed in light gray (onset at 0 ms). The colors of the OPM traces match the corresponding colors on the topographic plot in panel c. Thicker traces correspond to channels showing significant responses. Periods of statistically significant responses (i.e., exceeding the 95th percentile of the surrogate distribution) are denoted with a red line above. (**c**) Topographic plot of the sensors radial axes based on their value at the moment of highest activity (determined as the maximum of the root-mean-square across all sensors, 261 ms). Sensors showing significant responses are marked with red asterisks. (**d**) Source activity, reconstructed from the group-averaged response at its moment of highest activity (261 ms), displayed on a template MRI of a one-month-old infant, in the neurological convention. Only statistically significant sources are displayed.

### Oddball responses

In a second paradigm, we used a frequency tagging approach (Kabdebon et al., 2022; Regan, 1982) to demonstrate that our OPM-MEG setup can characterize auditory change detection in newborns. For that purpose, a sequence of pure tones was presented at a steady 3 Hz rate in trains of 3 standard tones followed by one oddball tone (Fig. 3a). The SNR of the radial field power spectra, averaged across the participants, showed clear peaks at the base stimulation frequency (3 Hz), the oddball frequency (0.75 Hz), and its first harmonic (1.5 Hz) (Fig. 3b), suggesting the presence of steady-state responses at the base and oddball frequencies. Some heartbeat artifacts remained after preprocessing, leading to SNR peaks in the 2.2-2.5 Hz range, interfering with the second oddball harmonic (2.25 Hz). We performed statistical tests at the frequencies of interest (0.75 Hz, 1.5 Hz and 3 Hz) and found that SNR peaks were all significant for the group-averaged response (*p* = 0.0016, *p* = 0.042, and *p* < 0.0001; maximum statistics), and that individual responses were significant in 1, 2 and 9 participants, respectively. The topographies for the group-averaged responses at 0.75 Hz, 1.5 Hz and 3 Hz showed that the corresponding SNR was localized above the temporal regions. The source localization showed activity peaking at the auditory cortices bilaterally at 0.75 Hz and 3 Hz, with a right-hemisphere dominance (Fig. 3b), though at 0.75 Hz significant activity in the left hemisphere is difficult to see because of its limited size.

**Fig. 3.**
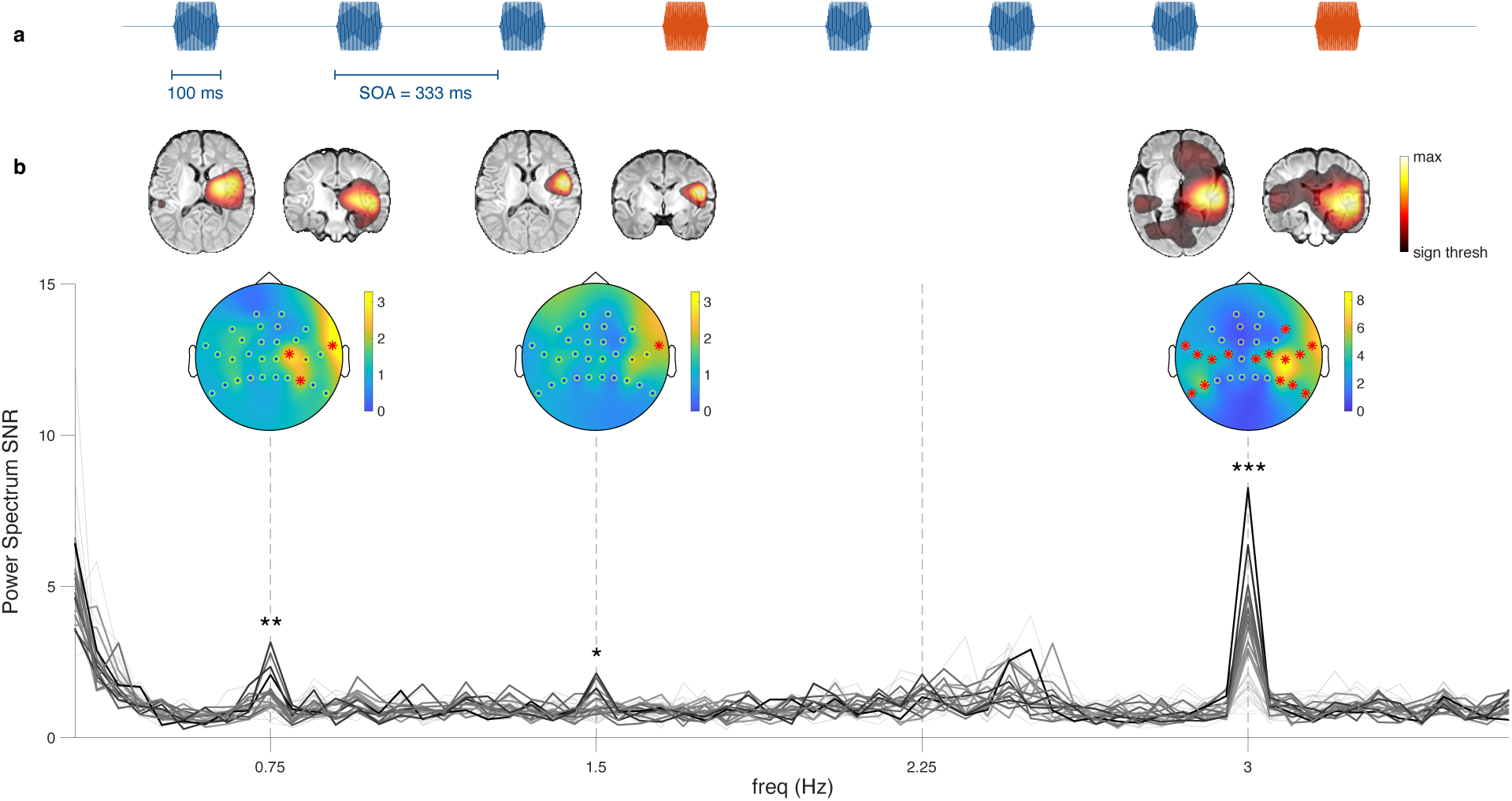
Oddball paradigm. (**a**) Excerpt from the sound sequence consisting of trains of 3 base tones (blue) followed by one oddball tone (orange). The base frequency is 3 Hz, and the oddball frequency is 0.75 Hz. (**b**) Group-averaged power spectrum SNR of the OPM radial axes; the display range is 0.3-3.6 Hz. Thicker lines represent channels that were found to be significant at one of the frequencies of interest. The shade of the individual sensor curves is obtained from their SNR value at 3 Hz. Topographic plots of the SNR at the corresponding frequency are displayed above the frequencies of interest. Channels with a significant SNR are marked by red asterisks. Above are shown the source localizations at those frequencies, on a template MRI of a one-month-old infant (brain images displayed in the neurological convention). Note that this is not the source projection of the topographic plots below (see Methods section). Only statistically significant sources are displayed.

### Hemispheric lateralization of auditory responses

The sources localizations of the responses enabled us to perform source level hemispheric comparisons, for which we used the maximally activated sources within the left and right hemisphere auditory cortices for the evoked response paradigm (Fig. 2d). The corresponding evoked responses are shown in Fig. 4a. Neither the amplitude nor the latency differed significantly between left and right auditory cortices (Fig. 4b-c). For the oddball paradigm, the bilaterality with right-hemisphere dominance of the power spectrum SNR was confirmed by plotting the power spectrum SNR of the left and right auditory sources (Fig. 4d). While some participants had higher SNRs in the right hemisphere, the statistical comparisons across individuals turned out to be not significant, for each of the frequencies of interest (Fig. 4e-g).

**Fig. 4.**
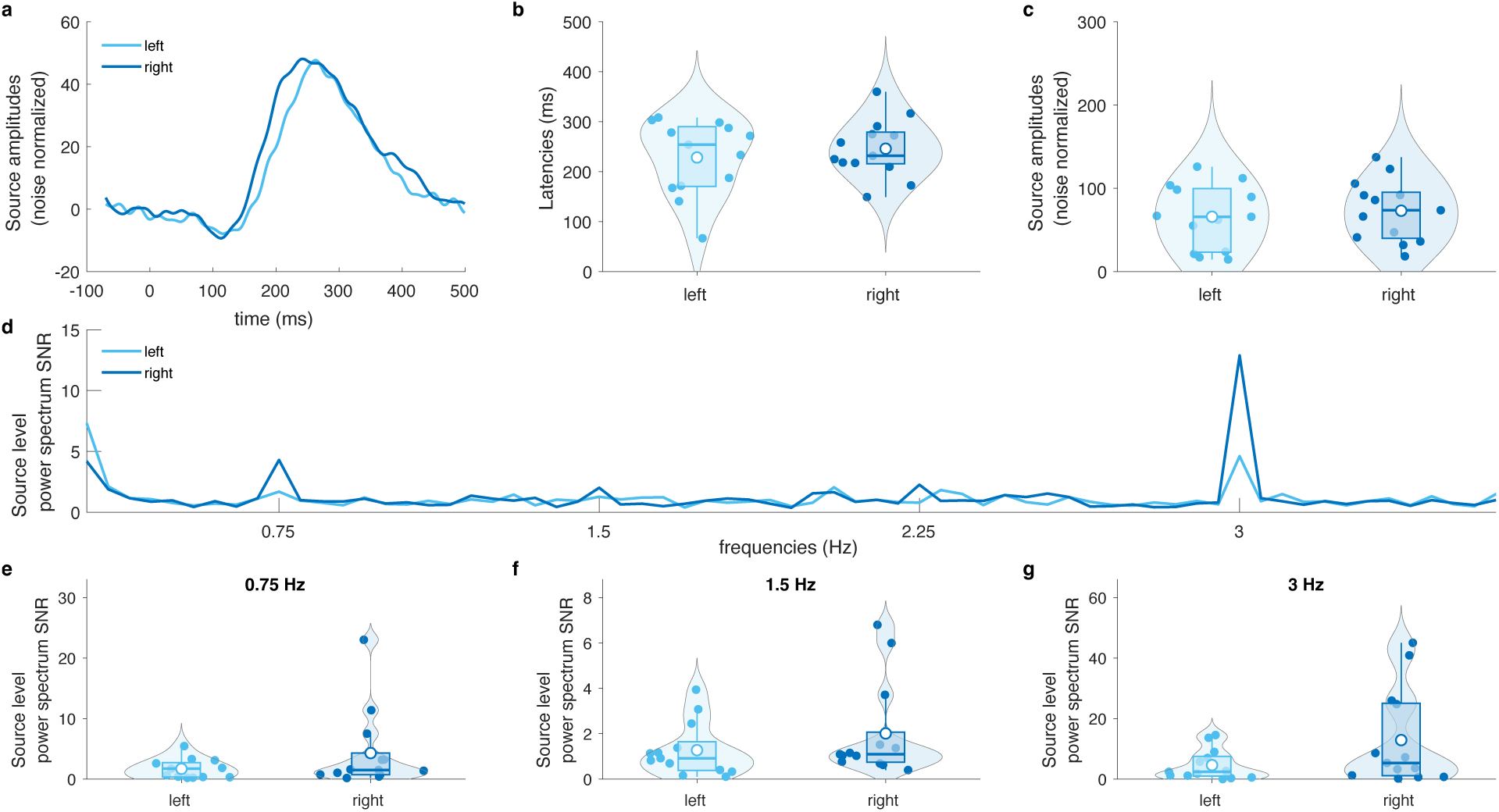
Hemispheric lateralization of auditory responses, in the evoked response paradigm (a-c) and in the oddball paradigm (d-g). (**a**) Auditory evoked responses at the maximally activated sources in the left (light blue) and right (dark blue) hemisphere. For all the remaining analyses of this figure, we used the two same sources and the reconstructions were made with the same inverse operator. (**b**) Comparison of the left and right responses latencies across the subjects, obtained as the moment where the responses peak. (**c**) Comparison of the left and right response peak amplitudes across the subjects. (**d**) Source level power spectrum SNR for the left and right sources. (**e**)-(**g**) Comparisons of the left and right source-level power spectrum SNR across the subjects at the frequencies of interest (0.75Hz, 1.5Hz, 3Hz). None of these comparisons were statistically significant.

### OPM tangential axes

For 10 out of the 14 participants, ∼20 triaxial OPMs (Gen3 QZFM, QuSpin Inc.) were available and placed over the temporal regions. In the first paradigm, the tangential component of the fields (normed and averaged across participants) displayed a clear response to the stimuli (Fig. 5a). This response peaked at 258 ms and the associated topographic plot exhibited peak activity directly above the dipole source (Fig. 5b), i.e., between the two poles observed with the radial axes (Fig. 2c). This indicates that both radial and tangential axes recorded the same activity. The response was statistically significant from 110 ms to 364 ms poststimulus onset (*p* < 0.0001, maximum statistics). Individual-level analyses revealed significant responses in 7 participants. In the second paradigm, the norm of the SNR of the OPM tangential power spectra, averaged across participants, exhibited peaks at the frequencies of interest (0.75 Hz, 1.5 Hz and 3 Hz; Fig. 5c), and all peaks were significant (*p* = 0.0012, *p* = 0.0007, *p* < 0.0001, maximum statistics). At the individual level, the responses were significant in 3, 3 and 6 participants respectively. We then compared the tangential and radial axes responses by evaluating the individual amplitudes and SNRs of the evoked responses and the power spectrum SNRs at the base and oddball frequencies (Fig. S2 in the Supplemental Material). The power spectrum SNR at 0.75 Hz was significantly higher for the tangential axes than for the radial axes (*W* = 7, Z = −2.09, *p* = 0.037, Wilcoxon signed-rank test). No other comparison was significant (all *p* > 0.1). The correlation between the amplitudes of the radial and tangential axes responses was very strong (*r* = 0.9, *p* = 0.0003).

**Fig. 5.**
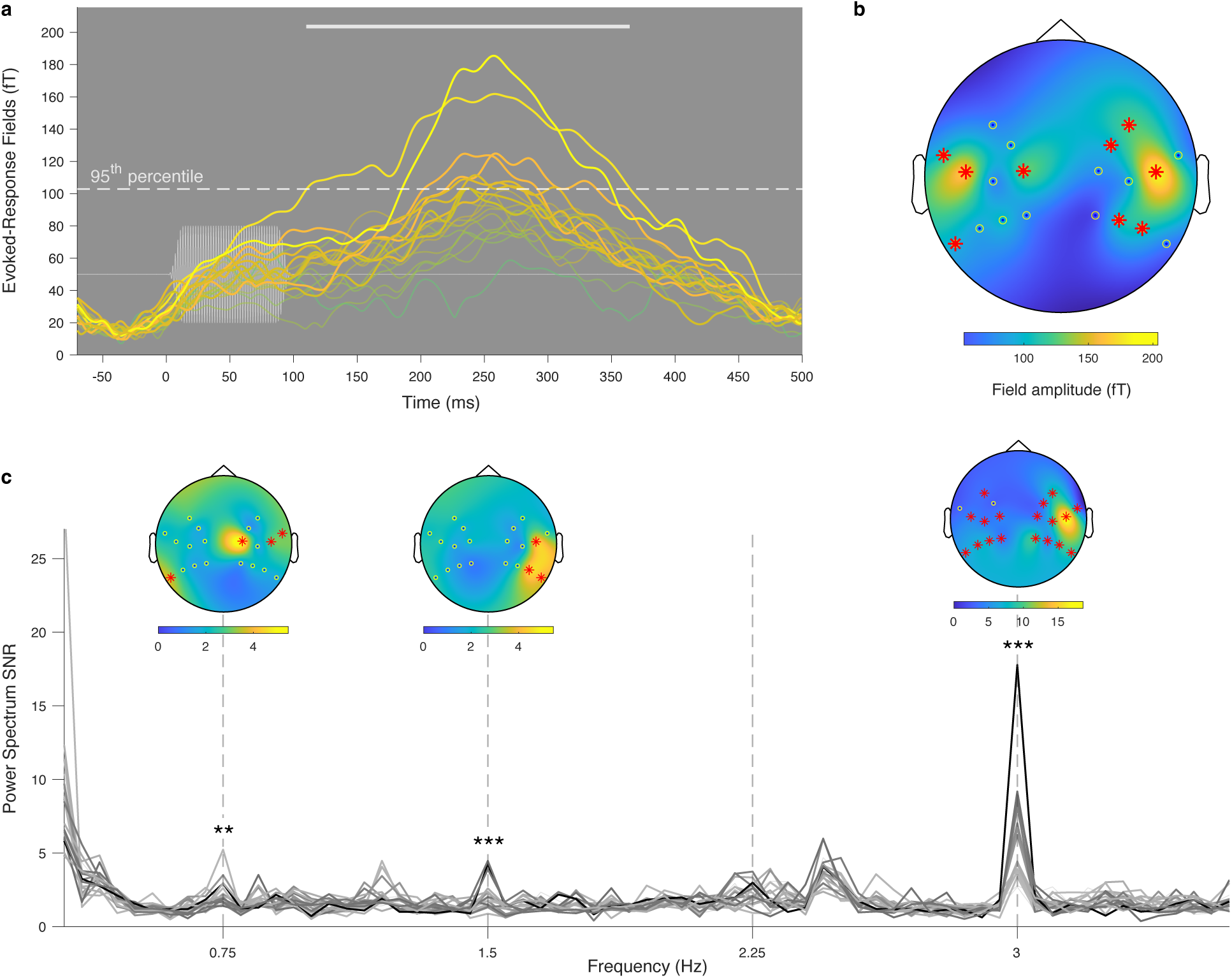
Responses in the direction tangential to the scalp for the 10 participants that had triaxial OPMs placed over their temporal regions. (**a**) Norm of the tangential magnetic field responses to pure tones and (**b**) the associated topography at 258 ms. Note that the response amplitude peaks right above the source currents that can be inferred based on Fig. 2c. This is consistent with the fact that cortical sources yield magnetic fields that are mostly radial to the scalp in their neighborhood and tangential to the scalp right above the source. The rest of the plot is the same as in Fig. 2. (**c**) Norm of the tangential power spectrum SNRs in the oddball steady-state experiment. The rest of the plot is the same as in Fig. 3.

### Comparison to cryogenic MEG

A subset of 7 infants were evaluated with the same paradigms in an adult-size cryogenic MEG system in the supine position, with their head laid sideways on the posterior sensors region. Our goal was to compare the temporal and spectral dynamics of auditory evoked and steady-state neural responses obtained using both modalities. The root-mean-square of the average auditory evoked responses peaked at 264 ms poststimulus onset with the OPM-MEG and at 282 ms with the cryogenic MEG. We compared the amplitudes and SNRs of the evoked responses and the power spectrum SNRs at the base and oddball frequencies (Fig. S3 in the Supplemental Material). All these comparisons were done at the sensors level. The amplitude of the evoked responses was significantly higher (on average 4 times higher) with OPM-MEG than with cryogenic MEG (*t*(6) = 8.05, *p* = 0.0002, *t-*test). This was to be expected due to the reduced brain-to-sensor distance (see e.g. (Zahran et al., 2022) for simulations). The SNRs of both the evoked responses and the power spectrum were not significantly different (all *p* > 0.26).

## Discussion

In this work, we developed a wearable OPM-MEG setup that fits and is well tolerated by newborns. Our design, which used flexible EEG-like caps and slightly elevated OPM holders, prevented heat-related discomfort to the infants’ skin. We demonstrated that this close-to-scalp OPM-MEG system is able to acquire high-quality data by performing two auditory stimulation paradigms in a group of 14 one-month-old healthy newborns.

In the first paradigm, the system successfully and accurately recorded the response evoked by pure tones. Its main component peaked at 261 ms, fully in agreement with the dominant P2 component previously described at that age in the EEG (Barnet et al., 1975b; Fellman & Huotilainen, 2006; Shibasaki & Miyazaki, 1992; Wunderlich et al., 2006) and cryogenic MEG (Draganova et al., 2007; Holst et al., 2005; Huotilainen et al., 2003; Lengle et al., 2001) literatures. In the second paradigm, we demonstrated the possibility of using our OPM-MEG system to perform frequency tagging paradigms with newborns, which have been shown to be very fruitful with young participants (Bertels et al., 2020; Kabdebon et al., 2022). While frequency tagging paradigms related to the one applied in this study were carried out with EEG in adults (Nozaradan et al., 2017) and infants (Cirelli et al., 2016; Edalati et al., 2023; Lenc et al., 2023), steady-state responses have not yet been studied in infants with MEG, nor with tone frequency changes. The power spectrum SNR exhibited significant peaks at the base frequency, at the oddball frequency and at its first harmonic. The last two are a frequency tagging analog of the auditory mismatch negativity (MMN), reflecting automatic change detection (Näätänen et al., 2007). In the frequency tagging approach, these peaks can also reveal rhythm perception and, more generally extraction of structural patterns (Cirelli et al., 2016; Moser et al., 2020; Lenc et al., 2023; Edalati et al., 2023), as the trains of stimuli are presented periodically. It is worth noting that observing an SNR peak in the spectrum at 0.75 Hz is a nontrivial feat of our system, as the OPM signal is known to be very noisy at such low frequencies due to movement and ambient field drifts (Boto et al., 2018). Dealing with such low-frequency noise has rarely been performed (de Lange et al., 2021). Hence, this OPM-MEG setup not only enabled us to evidence responses evoked by pure tones in the newborn brain, but also its ability to detect sound changes and regular sequences. In future studies, we should carefully consider the behavioral state of the infants tested. Previous studies have indeed shown that it can influence the amplitude of brain responses to changes in tone sequences (Duclaux et al., 1991; Moser et al., 2020).

Performing sources reconstruction, we were able to localize the sources of scalp magnetic fields within the infant brain at the auditory cortices, as expected. The source-level responses did not show any significant hemispheric lateralization, either in terms of latency or amplitude. The absence of hemispheric differences in response latency echoes previous findings (Wunderlich et al., 2006). Nevertheless, it is possible that an earlier response in the right hemisphere, as found in several studies, see, e.g., (Y. Chen et al., 2023; Musacchia et al., 2013), does not come out significant in our analysis due to the small sample size. Studies investigating the hemispheric differences in auditory response amplitudes are not in unison, with some authors reporting no hemispheric preference as we do (Y. Chen et al., 2023; Wunderlich et al., 2006), while others report higher electric potential over the right (Mento et al., 2010; Molfese et al., 1975; Musacchia et al., 2013) or left (Dehaene-Lambertz, 2000) hemisphere.

These source localization results were obtained from the group-averaged data, using a template MRI and an approximate estimation of sensors position and orientation with respect to that template. The fact that this coarse approach nonetheless led to an accurate source localization is a manifestation of the substantial advantage of the MEG signal not being distorted by the head tissues, compared to the EEG signal. In particular for the population of interest, the EEG signal is distorted due to sutures and fontanels (Lew et al., 2013), and individual realistic head models are necessary. Considering the difficulty of obtaining structural MRI in infants (Aeby et al., 2013; Wang et al., 2019), which largely impacts the recruitment of participants, the proposed approach demonstrates the key added value of OPM-MEG to perform source localization based on a template MRI in this population. In the near future, the advent of fast (<1 min) and precise (∼1 mm) automatic OPM co-registration (localization and orientation) on the subjects’ scalp using external coils, such as the zero field nulling coils of the MSR or a halo of fixed head positioning coils, will improve the accuracy of source localization (Hill et al., 2024; Livanainen et al., 2022; Pfeiffer et al., 2018, 2020).

Additionally, we showed that the neural activity is also accurately recorded by the OPMs in the direction tangential to the scalp, meaning that triaxial OPMs are a great way to increase the channel count on small heads without impacting the size and weight of the device. Although in our setup the activities captured in the radial and tangential directions were similar, as seen from the high correlation between the response amplitudes, they turned out to be better suited to reveal significant responses in the frequency tagging paradigm. In scenarios with even smaller head sizes, such as preterm neonates, the risk of missing activity with the radial axes alone becomes higher, and having tangential axes increases the chances that all the relevant activity is captured.

Lastly, we validated our results with an adult-size cryogenic MEG. The latencies of the responses were concordant, and the amplitudes significantly higher for the OPM-MEG, as expected given that the sensors are closer to the scalp in our setup. Nevertheless, we found similar SNRs for the two MEG types. Indeed, the noise level of the OPM signal is higher, which originates in movements creating high-amplitude artifacts, due to the sensors being displaced in a nonzero magnetic field. This issue can be tempered by keeping the ambient field as low as possible, which requires strong magnetic shielding and the use of field-nulling coils, as applied in our study. Further improvements in interference suppression techniques (Holmes, Bowtell, et al., 2023; Seymour et al., 2022) will likely increase the SNR of OPM data, which may ultimately exceed that of cryogenic MEG data. Another limitation of the comparisons of the evoked responses and their SNR performed here is that they were carried out at the sensors level, as we did not perform source localization for the cryogenic MEG. Further studies comparing modalities, benefiting from automatic OPM localization, should include continuous head localization using HPI coils for the cryogenic MEG, enabling sources level comparisons.

Compared to cryogenic MEG, OPM-MEG permits close-to-scalp recordings in infants without the need to build an expensive dedicated system. It is also much less energy intensive. Importantly, in our setup, data can be acquired in a setting that is naturalistic and appropriate for newborns, with the baby resting comfortably on their parent’s lap. This option could be even more valuable when testing older infants and toddlers, who may be more distressed than their younger peers by being physically separated from their caregiver. While a lot of efforts have been made to perform cryogenic MEG recordings with source localization in infants (see, e.g., (Y.-H. Chen et al., 2019; Clarke, Bosseler, et al., 2022; Clarke, Larson, et al., 2022; Kao & Zhang, 2019; Lew et al., 2013), cryogenic MEG will not be able to achieve this naturalness of recording. However, it is worth stressing that cryogenic MEG is a mature technology whereas OPM-MEG is still in its early stages.

Similarly, this level of naturalness is likely unachievable with rigid wearable helmets used in previous OPM studies (Hill et al., 2019; Rier et al., 2024). Our OPM-MEG setup offers the flexibility and applicability to newborns of wearable systems such as EEG, whilst providing the high spatial precision of MEG (for combination and differences of both modalities, see (Boto et al., 2019)). Nevertheless, as with EEG and fNIRS, children must agree to wear a cap, which can be problematic notably in certain populations of children. This might especially be the case in children with atypical development (e.g., autism spectrum disorders). For these populations, it might be easier to use a rigid helmet as is done in cryogenic MEG.

Our setup can also be easily adapted to older participants (for studies in school-aged children, see (Feys, Corvilain, Van Hecke, et al., 2023; Feys et al., 2022; Rier et al., 2024), making it applicable across the whole lifespan. Such experimental configuration opens the door to investigations of parent-child interactions in natural conditions, including hyperscanning (Holmes, Rea, Hill, Boto, et al., 2023; Nguyen et al., 2020). For toddlers and young children, one can envisage experiments where the participants are walking around or playing games. Indeed, in adults, successful OPM-MEG brain recordings have been achieved while participants were undergoing large movements (Holmes, Rea, Hill, Leggett, et al., 2023; Mellor et al., 2022, 2023), standing (Seymour et al., 2021), playing ping-pong (Boto et al., 2019; Holmes, Rea, Hill, Boto, et al., 2023) and guitar (Hill et al., 2019). It should be noted, however, that scanning older infants or toddlers may lead to new challenges to be addressed, as they are more likely to touch or manipulate the OPMs or their cables, or try to remove the cap. Generally speaking, the future of neurodevelopmental research likely resides in multimodal neuroimaging in which each modality, be it fMRI, EEG, fNIRS, cryogenic MEG or OPM-MEG, should be chosen according to the research question, the brain activity targeted and the feasibility.

In conclusion, the present study reports an innovative wearable MEG system that allows for more naturalistic recordings in neonates. This research paves the way for further early neurodevelopmental investigations using OPM-MEG, which will benefit from the increased naturalness, signal amplitude and spatial resolution from the close-to-scalp sensors. This may ultimately provide a wealth of unprecedented information on brain functional maturation from the very early stages of life.

## Materials and Methods

### Participants

Fourteen neonates (7 females) participated in this study; their ages ranged between 29 and 36 days (mean: 32.6 days). The inclusion criteria were as follows: gestational age >36 weeks, APGAR scores >7 at 5 and 10 minutes after birth, uncomplicated pregnancy, no known developmental delay, and normal hearing at birth (cf. otoacoustic emissions). For each paradigm, we rejected the data of one participant (a different one in each dataset) due to technical issues.

### System design and data acquisition

To build a wearable OPM-MEG system that was suitable for newborns, we used flexible and slightly stretchy caps (EasyCap, sizes: 36, 38 and 40 cm of head circumference), onto which we sewed OPM holders that we designed and 3D-printed with acrylonitrile butadiene styrene (ABS) on site. As a result, the interior of the cap was fabric, and the cap was comfortable to wear. The holders were set so that a gap of 4 mm was left between the scalp and the OPM (Fig. 1b), which is crucial to dissipate the heat generated by the rubidium-based OPMs and prevent hurting the sensitive skin of newborns. Indeed, the rubidium inside the sensitive cell needs to be heated to approximately 150°C to work in the spin exchange relaxation-free regime (Tierney et al., 2019), making the OPM surface relatively hot. Notably, helium-based OPMs do not heat (Gutteling et al., 2023); however, to date, He-OPMs are too heavy and bulky (Feys, Corvilain, Labyt, et al., 2023) to be placed on a newborn scalp, as opposed to the OPMs used in this study (size: 1.2 x 1.7 x 2.6 cm^3^, weight: 4.5–4.7 g). In addition, we attached aerogel foam to the bottom of the OPMs to further isolate the scalp. The caps were equipped with 19 to 30 (mean: 27) biaxial or triaxial OPMs (Gen2/Gen3 QZFM, QuSpin Inc.), depending on the cap size and OPM availability. The 15 to 24 triaxial sensors (mean: 20, available for 10 participants) were placed in priority over the temporal areas to increase sensitivity to auditory responses (Brookes et al., 2021). When wearing the cap, the newborns could rest on the lap of one of their parents, seated inside a magnetically shielded room (Compact MuRoom, MSL Ltd). The newborns were in a quiet state with eyes closed, and they all seemed to have fallen asleep. The room was degaussed and wall coils (MuCoils, Cerca Magnetics) were used—available for 10 participants—in order to further reduce the remnant magnetic fields below 1 nT within a 40-cm wide cubic region of interest containing the newborn’s head. This was achieved by mapping, before the acquisition, the remnant static magnetic field in that cube using the offset-corrected field-zero values of ∼20 triaxial OPMs positioned in that region using a custom-made all-wood structure. The magnetic profiles of the coils were determined at those positions by sending voltages in step pulses and measuring the magnetic responses with the OPMs. The latter were then inverted to null the magnetic field measured in the previous step. Data were recorded at a sampling frequency of 1200 Hz using a 16-bit digital acquisition (DAQ) system (National Instruments) controlled by a custom-made Python program. Audio stimuli were recorded simultaneously with the DAQ. If needed, the infants used a nonmagnetic pacifier. Live video was available during the whole experiment to monitor and address any problems that may have occurred. ***Cryogenic MEG.*** A subset of 7 subjects underwent the same paradigms in an adult-size TRIUX Neuromag (MEGIN) cryogenic MEG system with 306 channels (of which 102 are magnetometers) housed in a different magnetically shielded room (Maxshield, MEGIN; lightweight version with internal active feedback compensation system) than the OPM-MEG system. To maximize the subjects’ comfort and cooperation, and thus our chances of keeping them in a quiet state, no head position tracking method was used for these recordings, reducing the length of the preparation (no placement of head position indicator coils, no digitization of these coils). The subjects had their left hemisphere placed over the bottom of the cryogenic sensors array positioned in the supine position, ending up with their left temporal region being in front of the occipital sensors for a regular use with an adult. The recordings were performed in this single position, as the full procedure (OPM-MEG + cryogenic MEG) was already quite lengthy. Indeed, both OPM-MEG and cryogenic MEG recordings were performed on the same day. Their order was balanced within the flexibility we had given the availability of both systems. ***Direct energy consumption***. The OPMs (QuSpin Inc.) have a total direct power consumption of 5 W per sensor (*QZFM Gen-3 – QuSpin*, n.d.) and can be switched off when not in use. A high-density array of 128 OPMs (384 channels) used for 7 hours in one day would thus consume 4.4 kWh of electricity. For comparison, the 306-channel cryogenic MEG TRIUX neo (MEGIN) system has a minimum power consumption of 4.2 kW (*MEGIN-TRIUX-Neo-Product-Data-2021.Pdf*, n.d.) and cannot be switched off. It therefore consumes a minimum of 100.8 kWh of electricity in one day. Taking the helium recycler (*MEGIN-Internal-Helium-Recycler-2021.Pdf*, n.d.) and the type of current used into account, the daily consumption ranges between 110 and 190 kWh. We report here only the direct energy consumption, as no life cycle analysis of both MEG types has been performed. The corresponding carbon dioxide emissions will depend on the carbon intensity of the local electricity production. However, to have an idea, one can look at the CO2-equivalent emitted assuming the mean carbon intensity of electricity production across the world, i.e., 438 gCO2e/kWh in 2022 (*“Data Page: Carbon Intensity of Electricity Generation”, Part of the Following Publication: Hannah Ritchie, Pablo Rosado and Max Roser (2023) - “Energy”. Data Adapted from Ember, Energy Institute. Retrieved from* Https://Ourworldindata.Org*/Grapher/Carbon-Intensity-Electricity [Online Resource]*, n.d.). In that scenario, the use of OPM-MEG would emit 0.7 tCO2e/year for 128 OPMs used 7 hours every single day, while possessing a cryogenic MEG would emit 17.6 to 30.4 tCO2e/year, regardless of its utilization.

### Auditory stimuli

Both auditory paradigms consisted of 5-minute sequences of pure tones, each lasting 100 ms including 15 ms of fade-in and fade-out, presented at 65 dB using a flat MEG-compatible speaker (Panphonics). The sequences were generated using a custom-made Python program (Python Language Reference, version 3.7). We presented each type of sequence in a randomized order and, when possible, more than once to increase the amount of data. Each subject had a minimum of 1 recording for each paradigm, with a maximum of 6 recordings when considering both paradigms. The mean total recording time was 20 ± 6 min for the OPM-MEG recordings and 16 ± 2 min for the cryogenic MEG recordings. The recording times per paradigm are presented in Table S1 in the Supplemental Material. ***Evoked response paradigm.*** In each sequence, a single pure tone with a frequency of either 500 Hz or 750 Hz (depending on the sequence) was presented with a stimulus onset asynchrony (SOA) randomly varying between 500 and 600 ms (Fig. 2a). ***Oddball paradigm.*** Pure tones were presented at a steady 3 Hz rate in trains of 3 base tones followed by one oddball tone, leading to an oddball frequency of 0.75 Hz (Fig. 3a); this paradigm was inspired by (Nozaradan et al., 2017). The base (oddball) tone frequency was either 500 or 750 Hz (750 or 500 Hz), depending on the sequence. For both paradigms, the sequences with different tone frequencies were treated identically for the analysis.

### Data analysis

All the data preprocessing and analyses were performed with custom-made scripts for Matlab (MathWork Inc., version 2020a-2024a), unless otherwise stated. Topographic plots were generated using the Fieldtrip toolbox (Oostenveld et al., 2011). ***Preprocessing*.** Recording periods containing large movement artifacts were first identified by visual inspection of the continuous data and marked as artifact time windows. Heartbeat artifacts from the newborn and the parent were then removed with an independent component analysis (ICA) (Vigário et al., 2000) using the FastICA algorithm (Hyvarinen, 1999), which was applied to the artifact-free part of the recordings, bandpass filtered between 1-40 Hz. Time windows affected by remaining artifacts were identified by visual examination of the other independent components by sliding through 5-second windows. The amount of data identified as contaminated by artifacts at the various analysis stages is described in Table S1 in the Supplemental Material, both for the OPM-MEG and cryogenic MEG data. ***Evoked response paradigm***. To avoid filtering artifacts, data in the artifact periods were replaced by linear interpolation before being bandpass filtered between 1-40 Hz and notch filtered at the frequencies at which the power spectrum displayed noise peaks, within the 13-40 Hz range. Artifacts periods were extended by 1.5 s in both directions. A robust z-score rejection (channel per channel, threshold value of 5, data amplitude smoothed with a 1 sec square window beforehand) was performed to identify samples affected by remaining artifacts. The first three principal components of the artifact-free data were projected out (Feys et al., 2022). The resulting data were decomposed into epochs from −70 to 500 ms around the start of each sound presentation (trial), and epochs that intersected artifact periods were rejected. The amount of recorded and accepted trials is presented in Table S2 in the Supplemental Material. The remaining epochs were baseline corrected, with a baseline spanning the 70 msprestimulus period. The epochs were averaged for each participant (individual-level evoked response). The norm of the response along the tangential direction was taken. Finally, the evoked responses were averaged across all participants (group-level). ***Oddball paradigm*.** The data were bandpass filtered between 0.2-40 Hz (to maintain access to the oddball frequency), and notch filtered at the frequencies at which the power spectrum displayed noise peaks, within the 13-40 Hz range. The first three principal components estimated from the artifact-free data were projected out. The ranges of the newborn’s and parent’s heart rates were computed based on the components identified in the ICA preprocessing, and further components containing heartbeat artifacts were removed if, in those heart frequency ranges, their power spectrum SNR (computed as explained below) exceeded 7.5. The resulting data were decomposed into artifact-free, 20-s epochs, starting when an oddball sound was presented to preserve the stimulus-related temporal structure across epochs. The amount of recorded and accepted 20-s epochs is presented in Table S2. The complex-valued Fourier coefficients of the data in each epoch were computed using the fast Fourier transform algorithm and averaged across epochs. The power spectrum was obtained as the squared magnitude of the resulting Fourier coefficients. Then, the SNR was computed for each frequency bin as the ratio of the power at that frequency to the mean power across the two 0.1-0.4 Hz frequency intervals around the considered frequency. The tangential SNR was computed as the norm of the SNRs in the 2 tangential directions. Group-level SNRs were obtained by averaging across participants. ***Cryogenic MEG***. Data were first denoised using signal space separation (MaxFilter, MEGIN; including subtraction of artefacts generated by internal active compensation) without movement compensation, as the head position was not tracked. Only the magnetometer data were retained, and all the following steps were the same as those for processing the OPM-MEG data, including the artifacts removal by visual inspection and the ICA to remove the heart component. The only difference was that the 3 first principal components were not removed since signal space separation was already applied. The lengths of recorded data, of artifact-free data, and the number of accepted trials did not statistically differ between the two MEG systems (see Fig. S4 in the Supplemental Material). ***Group averages*.** Classical group averaging of sensors data was used to summarize the common response pattern across subjects. These averages were performed based on a 2D template layout that most closely matched the sensors positions for all the different cap sizes used (custom-made for OPM-MEG, using the Neuromag (MEGIN) layout implemented in Fieldtrip for cryogenic MEG). Note that this procedure of averaging at the sensors level inevitably yields topographic plots that are smoother and less focal than the individual ones***. Source localization*.** In the absence of MRIs of the individual participants, source generators of OPM responses were localized within a template MRI of a one-month-old infant, sourced from the Brain Connectome Project (BCP) (L. Chen et al., 2022). To register the OPMs positions and orientations on this template, the head surface of the smallest available Montreal Neurological Institute (MNI) template (Fonov et al., 2009) was used, as the BCP image is devoid of skin information. This head surface was rescaled to match the head circumference of our group average and centered on the BCP MRI brain volume. It was then 3D printed, the flexible cap mounted on it and populated with OPMs. The positions and orientations of the OPMs, together with the surface of the 3D printed face, were recorded using a 3D scanner (Einscan H2). Co-registration with the template MRI was performed using Blender with a co-registration addon (Zetter et al., 2019), see Fig. S1. This co-registration, *a posteriori* and on a template MRI, could not be achieved for the cryogenic system, so that source localization was only performed for OPM-MEG. The forward model was computed using the one-layer Boundary Element Method of the MNE software (Gramfort et al., 2014). It is worth noting that the MEG forward model is rather insensitive to head modeling approximations, as opposed to EEG, in particular for the skull bone shape (Baillet, 2017; Lanfer et al., 2012). With this in mind, the inner skull surface used in this model was obtained from the adult MNI inner skull surface, shrunk until it fitted snugly to the one-month-old BCP brain volume, as the latter could not be segmented. Using the Minimum-Norm Estimates inverse solution, with depth bias correction by noise normalization (Dale & Sereno, 1993), sources activations were reconstructed for the peak activity of the group averaged evoked response. For the oddball paradigm, sources cannot be reconstructed from group-averaged SNR as this quantity is not linear in the magnetic field. Instead, they were reconstructed per subject for the Fourier coefficients at the frequencies of interest. The power spectra and their SNRs were then computed, and the group-average was taken. For all source localizations, all OPMs available across the whole group were taken into account (30 in total, 15 per hemisphere) but restricted to their radial axis.

### Statistics

Maximum statistics tests were performed on the evoked and steady-state responses at both the individual and group levels, hence providing full control of the familywise error rate. The iterative *step-down method* (Nichols & Holmes, 2002) was used, i.e., data that were found to be significant in one iteration were masked off the maximum statistics test in the next iteration until no more significant data were found. The statistical significance level was set to *p* < 0.05. Null distributions were constructed from 10,000 samples of surrogate data built according to the null hypothesis that they contain no response to the stimuli. For the evoked responses in the time domain, the sign of each epoch was randomly flipped (permutation test), and for the frequency tagging analysis, the phases of Fourier coefficients were randomly changed. In both cases, the same changes were applied to all OPM channels to preserve their spatial distribution. To compare the different modalities (radial vs. tangential axes, n = 10, and OPM vs. cryogenic MEG radial magnetometers, n = 7) and to evaluate hemispheric lateralization (n = 13) we performed two-tailed paired sample *t*-tests, unless Shapiro‒Wilk tests indicated that the distributions deviated from normality, in which case two-tailed Wilcoxon signed-rank tests were performed. To compare the full systems (tri-axial OPMs vs. magnetometers and gradiometers of cryogenic MEG, n=6), we performed two-tailed unpaired sample *t*-tests, unless Shapiro‒Wilk tests indicated that the distributions deviated from normality, in which case two-tailed Wilcoxon rank-sum tests were performed. For these tests, the amplitudes of the evoked responses were computed as the highest value across the channels at the point where the principal component of the response peaked. The SNR for these evoked responses was computed as the ratio between the amplitude and the standard deviation of the corresponding channel time-series during the baseline (70 ms prestimulus period). Finally, the individual power spectrum SNRs for the steady-state responses were taken as the highest power spectrum SNR value at each of the frequencies of interest.

## Additional Information

### Author contributions

Conceptualization: PC, JB, VW, XDT

Methodology: PC

Software: PC

Formal analysis: PC, VW, MB

Investigation: PC, JB, CC, LF, MF

Resources: XDT, JB, CC, OF

Data Curation: PC

Writing—original draft: PC

Writing—review & editing: PC, JB, VW, MB, XDT, CC, MF, OF, LF

Visualization: PC

Supervision: JB, XDT

Funding acquisition: JB, XDT

### Ethics

The parents gave informed consent prior to testing and received monetary compensation for their participation. The CUB Hôpital Erasme Ethics Committee approved the experimental protocol (P2021/141/B406202100081). The methods were carried out in accordance with the approved guidelines and regulations.

### Data and materials availability

All data, code, and materials used in the analyses have been archived to an open data publishing platform Zenodo and will be made publicly available upon publication at https://doi.org/10.5281/zenodo.10213856 (dataset) and https://doi.org/10.5281/zenodo.10213974 (code).

## References

Aeby, A., De Tiège, X., Creuzil, M., David, P., Balériaux, D., Van Overmeire, B., Metens, T., & Van Bogaert, P. (2013). Language development at 2 years is correlated to brain microstructure in the left superior temporal gyrus at term equivalent age: A diffusion tensor imaging study. NeuroImage, 78, 145–151. 10.1016/j.neuroimage.2013.03.076

Baillet, S. (2017). Magnetoencephalography for brain electrophysiology and imaging. Nature Neuroscience, 20(3), 327–339. 10.1038/nn.4504

Barnet, A. B., Ohlrich, E. S., Weiss, I. P., & Shanks, B. (1975a). Auditory evoked potentials during sleep in normal children from ten days to three years of age. Electroencephalography and Clinical Neurophysiology, 39(1), 29–41. 10.1016/0013-4694(75)90124-8

Barnet, A. B., Ohlrich, E. S., Weiss, I. P., & Shanks, B. (1975b). Auditory evoked potentials during sleep in normal children from ten days to three years of age. Electroencephalography and Clinical Neurophysiology, 39(1), 29–41. 10.1016/0013-4694(75)90124-8

Bertels, J., Bourguignon, M., de Heering, A., Chetail, F., De Tiège, X., Cleeremans, A., & Destrebecqz, A. (2020). Snakes elicit specific neural responses in the human infant brain. Scientific Reports, 10(1), 7443. 10.1038/s41598-020-63619-y

Boto, E., Holmes, N., Leggett, J., Roberts, G., Shah, V., Meyer, S. S., Muñoz, L. D., Mullinger, K. J., Tierney, T. M., Bestmann, S., Barnes, G. R., Bowtell, R., & Brookes, M. J. (2018). Moving magnetoencephalography towards real-world applications with a wearable system. Nature, 555(7698), 657–661. 10.1038/nature26147

Boto, E., Meyer, S. S., Shah, V., Alem, O., Knappe, S., Kruger, P., Fromhold, T. M., Lim, M., Glover, P. M., Morris, P. G., Bowtell, R., Barnes, G. R., & Brookes, M. J. (2017). A new generation of magnetoencephalography: Room temperature measurements using optically-pumped magnetometers. NeuroImage, 149, 404–414. 10.1016/j.neuroimage.2017.01.034

Boto, E., Seedat, Z. A., Holmes, N., Leggett, J., Hill, R. M., Roberts, G., Shah, V., Fromhold, T. M., Mullinger, K. J., Tierney, T. M., Barnes, G. R., Bowtell, R., & Brookes, M. J. (2019). Wearable neuroimaging: Combining and contrasting magnetoencephalography and electroencephalography. NeuroImage, 201, 116099. 10.1016/j.neuroimage.2019.116099

Boto, E., Shah, V., Hill, R. M., Rhodes, N., Osborne, J., Doyle, C., Holmes, N., Rea, M., Leggett, J., Bowtell, R., & Brookes, M. J. (2022). Triaxial detection of the neuromagnetic field using optically-pumped magnetometry: Feasibility and application in children. NeuroImage, 252, 119027. 10.1016/j.neuroimage.2022.119027

Brookes, M. J., Boto, E., Rea, M., Shah, V., Osborne, J., Holmes, N., Hill, R. M., Leggett, J., Rhodes, N., & Bowtell, R. (2021). Theoretical advantages of a triaxial optically pumped magnetometer magnetoencephalography system. NeuroImage, 236, 118025. 10.1016/j.neuroimage.2021.118025

Brookes, M. J., Leggett, J., Rea, M., Hill, R. M., Holmes, N., Boto, E., & Bowtell, R. (2022). Magnetoencephalography with optically pumped magnetometers (OPM-MEG): The next generation of functional neuroimaging. Trends in Neurosciences, 45(8), 621–634. 10.1016/j.tins.2022.05.008

Chen, L., Wu, Z., Hu, D., Wang, Y., Zhao, F., Zhong, T., Lin, W., Wang, L., & Li, G. (2022). A 4D infant brain volumetric atlas based on the UNC/UMN baby connectome project (BCP) cohort. NeuroImage, 253, 119097. 10.1016/j.neuroimage.2022.119097

Chen, Y., Green, H. L., Putt, M. E., Allison, O., Kuschner, E. S., Kim, M., Blaskey, L., Mol, K., McNamee, M., Bloy, L., Liu, S., Huang, H., Roberts, T. P. L., & Edgar, J. C. (2023). Maturation of auditory cortex neural responses during infancy and toddlerhood. NeuroImage, 275, 120163. 10.1016/j.neuroimage.2023.120163

Chen, Y.-H., Saby, J., Kuschner, E., Gaetz, W., Edgar, J. C., & Roberts, T. P. L. (2019). Magnetoencephalography and the infant brain. NeuroImage, 189, 445–458. 10.1016/j.neuroimage.2019.01.059

Cirelli, L. K., Spinelli, C., Nozaradan, S., & Trainor, L. J. (2016). Measuring Neural Entrainment to Beat and Meter in Infants: Effects of Music Background. Frontiers in Neuroscience, 10, 229. 10.3389/fnins.2016.00229

Clarke, M. D., Bosseler, A. N., Mizrahi, J. C., Peterson, E. R., Larson, E., Meltzoff, A. N., Kuhl, P. K., & Taulu, S. (2022). Infant brain imaging using magnetoencephalography: Challenges, solutions, and best practices. Human Brain Mapping, 43(12), 3609–3619. 10.1002/hbm.25871

Clarke, M. D., Larson, E., Peterson, E. R., McCloy, D. R., Bosseler, A. N., & Taulu, S. (2022). Improving Localization Accuracy of Neural Sources by Pre-processing: Demonstration with Infant MEG Data. Frontiers in Neurology, 13, 827529. 10.3389/fneur.2022.827529

Copeland, A., Silver, E., Korja, R., Lehtola, S. J., Merisaari, H., Saukko, E., Sinisalo, S., Saunavaara, J., Lähdesmäki, T., Parkkola, R., Nolvi, S., Karlsson, L., Karlsson, H., & Tuulari, J. J. (2021). Infant and Child MRI: A Review of Scanning Procedures. Frontiers in Neuroscience, 15. 10.3389/fnins.2021.666020

Dale, A. M., & Sereno, M. I. (1993). Improved Localizadon of Cortical Activity by Combining EEG and MEG with MRI Cortical Surface Reconstruction: A Linear Approach. Journal of Cognitive Neuroscience, 5(2), 162–176. 10.1162/jocn.1993.5.2.162

“Data Page: Carbon intensity of electricity generation”, part of the following publication: Hannah Ritchie, Pablo Rosado and Max Roser (2023)—“Energy”. Data adapted from Ember, Energy Institute. *Retrieved from* https://ourworldindata.org/grapher/carbon-intensity-electricity *[online resource]*. (n.d.).

de Lange, P., Boto, E., Holmes, N., Hill, R. M., Bowtell, R., Wens, V., De Tiège, X., Brookes, M. J., & Bourguignon, M. (2021). Measuring the cortical tracking of speech with optically-pumped magnetometers. NeuroImage, 233, 117969. 10.1016/j.neuroimage.2021.117969

Dehaene-Lambertz, G. (2000). Cerebral specialization for speech and non-speech stimuli in infants. Journal of Cognitive Neuroscience, 12(3), 449–460. 10.1162/089892900562264

Draganova, R., Eswaran, H., Murphy, P., Lowery, C., & Preissl, H. (2007). Serial magnetoencephalographic study of fetal and newborn auditory discriminative evoked responses. Early Human Development, 83(3), 199–207. 10.1016/j.earlhumdev.2006.05.018

Duclaux, R., Challamel, M. J., Collet, L., Roullet-Solignac, I., & Revol, M. (1991). Hemispheric asymmetry of late auditory evoked response induced by pitch changes in infants: Influence of sleep stages. Brain Research, 566(1–2), 152–158. 10.1016/0006-8993(91)91693-u

Edalati, M., Wallois, F., Safaie, J., Ghostine, G., Kongolo, G., Trainor, L. J., & Moghimi, S. (2023). Rhythm in the Premature Neonate Brain: Very Early Processing of Auditory Beat and Meter. The Journal of Neuroscience: The Official Journal of the Society for Neuroscience, 43(15), 2794–2802. 10.1523/JNEUROSCI.1100-22.2023

Fellman, V., & Huotilainen, M. (2006). Cortical auditory event-related potentials in newborn infants. Seminars in Fetal & Neonatal Medicine, 11(6), 452–458. 10.1016/j.siny.2006.07.004

Feys, O., Corvilain, P., Aeby, A., Sculier, C., Holmes, N., Brookes, M., Goldman, S., Wens, V., & De Tiège, X. (2022). On-Scalp Optically Pumped Magnetometers versus Cryogenic Magnetoencephalography for Diagnostic Evaluation of Epilepsy in School-aged Children. Radiology, 304(2), 429–434. 10.1148/radiol.212453

Feys, O., Corvilain, P., Bertels, J., Sculier, C., Holmes, N., Brookes, M., Wens, V., & De Tiège, X. (2023). On-scalp magnetoencephalography for the diagnostic evaluation of epilepsy during infancy. Clinical Neurophysiology: Official Journal of the International Federation of Clinical Neurophysiology, 155, 29–31. 10.1016/j.clinph.2023.08.010

Feys, O., Corvilain, P., Labyt, E., Mahmoudzadeh, M., Routier, L., Sculier, C., Holmes, N., Brookes, M., Goldman, S., Romain, R., Mitryukovskiy, S., Palacios-Laloy, A., Schwartz, D., Betrouni, N., Derambure, P., Wallois, F., Wens, V., & De Tiège, X. (2023). Tri-axial rubidium and helium optically pumped magnetometers for on-scalp magnetoencephalography recording of interictal epileptiform discharges: A case study. Frontiers in Neuroscience, 17, 1284262. 10.3389/fnins.2023.1284262

Feys, O., Corvilain, P., Van Hecke, A., Sculier, C., Rikir, E., Legros, B., Gaspard, N., Leurquin-Sterk, G., Holmes, N., Brookes, M., Goldman, S., Wens, V., & De Tiège, X. (2023). Recording of Ictal Epileptic Activity Using on-Scalp Magnetoencephalography. Annals of Neurology, 93(2), 419–421. 10.1002/ana.26562

Feys, O., & De Tiège, X. (2024). From cryogenic to on-scalp magnetoencephalography for the evaluation of paediatric epilepsy. Developmental Medicine and Child Neurology, 66(3), 298–306. 10.1111/dmcn.15689

Feys, O., Ferez, M., Corvilain, P., Schuind, S., Rikir, E., Legros, B., Gaspard, N., Holmes, N., Brookes, M., Wens, V., & De Tiège, X. (2024). On-Scalp Magnetoencephalography Based On Optically Pumped Magnetometers Can Detect Mesial Temporal Lobe Epileptiform Discharges. Annals of Neurology, 95(3), 620–622. 10.1002/ana.26844

Fonov, V., Evans, A., McKinstry, R., Almli, C., & Collins, D. (2009). Unbiased nonlinear average age-appropriate brain templates from birth to adulthood. NeuroImage, 47, S102. 10.1016/S1053-8119(09)70884-5

Gramfort, A., Luessi, M., Larson, E., Engemann, D. A., Strohmeier, D., Brodbeck, C., Parkkonen, L., & Hämäläinen, M. S. (2014). MNE software for processing MEG and EEG data. NeuroImage, 86, 446–460. 10.1016/j.neuroimage.2013.10.027

Gutteling, T. P., Bonnefond, M., Clausner, T., Daligault, S., Romain, R., Mitryukovskiy, S., Fourcault, W., Josselin, V., Le Prado, M., Palacios-Laloy, A., Labyt, E., Jung, J., & Schwartz, D. (2023). A New Generation of OPM for High Dynamic and Large Bandwidth MEG: The 4He OPMs-First Applications in Healthy Volunteers. *Sensors (Basel*, Switzerland), 23(5), 2801. 10.3390/s23052801

Hill, R. M., Boto, E., Holmes, N., Hartley, C., Seedat, Z. A., Leggett, J., Roberts, G., Shah, V., Tierney, T. M., Woolrich, M. W., Stagg, C. J., Barnes, G. R., Bowtell, R., Slater, R., & Brookes, M. J. (2019). A tool for functional brain imaging with lifespan compliance. Nature Communications, 10(1), 4785. 10.1038/s41467-019-12486-x

Hill, R. M., Rivero, G. R., Tyler, A. J., Schofield, H., Doyle, C., Osborne, J., Bobela, D., Rier, L., Gibson, J., Tanner, Z., Boto, E., Bowtell, R., Brookes, M. J., Shah, V., & Holmes, N. (2024). *Determining sensor geometry and gain in a wearable MEG system* (No. arXiv:2410.08718). arXiv. 10.48550/arXiv.2410.08718

Holmes, N., Bowtell, R., Brookes, M. J., & Taulu, S. (2023). An Iterative Implementation of the Signal Space Separation Method for Magnetoencephalography Systems with Low Channel Counts. *Sensors (Basel*, Switzerland), 23(14), 6537. 10.3390/s23146537

Holmes, N., Rea, M., Hill, R. M., Boto, E., Leggett, J., Edwards, L. J., Rhodes, N., Shah, V., Osborne, J., Fromhold, T. M., Glover, P., Montague, P. R., Brookes, M. J., & Bowtell, R. (2023). Naturalistic Hyperscanning with Wearable Magnetoencephalography. *Sensors (Basel*, Switzerland), 23(12), 5454. 10.3390/s23125454

Holmes, N., Rea, M., Hill, R. M., Leggett, J., Edwards, L. J., Hobson, P. J., Boto, E., Tierney, T. M., Rier, L., Rivero, G. R., Shah, V., Osborne, J., Fromhold, T. M., Glover, P., Brookes, M. J., & Bowtell, R. (2023). Enabling ambulatory movement in wearable magnetoencephalography with matrix coil active magnetic shielding. NeuroImage, 274, 120157. 10.1016/j.neuroimage.2023.120157

Holst, M., Eswaran, H., Lowery, C., Murphy, P., Norton, J., & Preissl, H. (2005). Development of auditory evoked fields in human fetuses and newborns: A longitudinal MEG study. Clinical Neurophysiology: Official Journal of the International Federation of Clinical Neurophysiology, 116(8), 1949–1955. 10.1016/j.clinph.2005.04.008

Huotilainen, M., Kujala, A., Hotakainen, M., Shestakova, A., Kushnerenko, E., Parkkonen, L., Fellman, V., & Näätänen, R. (2003). Auditory magnetic responses of healthy newborns. Neuroreport, 14(14), 1871–1875. 10.1097/00001756-200310060-00023

Hyvarinen, A. (1999). Fast ICA for noisy data using Gaussian moments. 1999 IEEE International Symposium on Circuits and Systems (ISCAS), 5, 57–61 vol.5. 10.1109/ISCAS.1999.777510

Kabdebon, C., Fló, A., de Heering, A., & Aslin, R. (2022). The power of rhythms: How steady-state evoked responses reveal early neurocognitive development. NeuroImage, 254, 119150. 10.1016/j.neuroimage.2022.119150

Kao, C., & Zhang, Y. (2019). Magnetic Source Imaging and Infant MEG: Current Trends and Technical Advances. Brain Sciences, 9(8), 181. 10.3390/brainsci9080181

Kushnerenko, E., Ceponiene, R., Balan, P., Fellman, V., & Näätänen, R. (2002). Maturation of the auditory change detection response in infants: A longitudinal ERP study. NeuroReport, 13(15), 1843.

Lanfer, B., Scherg, M., Dannhauer, M., Knösche, T. R., Burger, M., & Wolters, C. H. (2012). Influences of skull segmentation inaccuracies on EEG source analysis. NeuroImage, 62(1), 418–431. 10.1016/j.neuroimage.2012.05.006

Lenc, T., Peter, V., Hooper, C., Keller, P. E., Burnham, D., & Nozaradan, S. (2023). Infants show enhanced neural responses to musical meter frequencies beyond low-level features. Developmental Science, 26(5), e13353. 10.1111/desc.13353

Lengle, J. M., Chen, M., & Wakai, R. T. (2001). Improved neuromagnetic detection of fetal and neonatal auditory evoked responses. Clinical Neurophysiology: Official Journal of the International Federation of Clinical Neurophysiology, 112(5), 785–792. 10.1016/s1388-2457(01)00532-6

Lew, S., Sliva, D. D., Choe, M., Grant, P. E., Okada, Y., Wolters, C. H., & Hämäläinen, M. S. (2013). Effects of sutures and fontanels on MEG and EEG source analysis in a realistic infant head model. NeuroImage, 76, 282–293. 10.1016/j.neuroimage.2013.03.017

Livanainen, J., Borna, A., Zetter, R., Carter, T. R., Stephen, J. M., McKay, J., Parkkonen, L., Taulu, S., & Schwindt, P. D. D. (2022). Calibration and Localization of Optically Pumped Magnetometers Using Electromagnetic Coils. Sensors, 22(8), Article 8. 10.3390/s22083059

Martynova, O., Kirjavainen, J., & Cheour, M. (2003). Mismatch negativity and late discriminative negativity in sleeping human newborns. Neuroscience Letters, 340(2), 75–78. 10.1016/s0304-3940(02)01401-5

MEGIN-Internal-Helium-Recycler-2021.pdf. (n.d.). Retrieved March 25, 2024, from https://megin.com/wp-content/uploads/2022/01/MEGIN-Internal-Helium-Recycler-2021.pdf

MEGIN-TRIUX-neo-Product-Data-2021.pdf. (n.d.). Retrieved March 25, 2024, from https://megin.com/wp-content/uploads/2022/01/MEGIN-TRIUX-neo-Product-Data-2021.pdf

Mellor, S., Tierney, T. M., OaNeill, G. C., Alexander, N., Seymour, R. A., Holmes, N., Lopez, J. D., Hill, R. M., Boto, E., Rea, M., Roberts, G., Leggett, J., Bowtell, R., Brookes, M. J., Maguire, E. A., Walker, M. C., & Barnes, G. R. (2022). Magnetic Field Mapping and Correction for Moving OP-MEG. IEEE Transactions on Bio-Medical Engineering, 69(2), 528–536. 10.1109/TBME.2021.3100770

Mellor, S., Tierney, T. M., Seymour, R. A., Timms, R. C., O’Neill, G. C., Alexander, N., Spedden, M. E., Payne, H., & Barnes, G. R. (2023). Real-time, model-based magnetic field correction for moving, wearable MEG. NeuroImage, 278, 120252. 10.1016/j.neuroimage.2023.120252

Mento, G., Suppiej, A., Altoè, G., & Bisiacchi, P. S. (2010). Functional hemispheric asymmetries in humans: Electrophysiological evidence from preterm infants. European Journal of Neuroscience, 31(3), 565–574. 10.1111/j.1460-9568.2010.07076.x

Molfese, D. L., Freeman, R. B., & Palermo, D. S. (1975). The ontogeny of brain lateralization for speech and nonspeech stimuli. Brain and Language, 2, 356–368. 10.1016/S0093-934X(75)80076-9

Moser, J., Schleger, F., Weiss, M., Sippel, K., Dehaene-Lambertz, G., & Preissl, H. (2020). Magnetoencephalographic signatures of hierarchical rule learning in newborns. Developmental Cognitive Neuroscience, 46, 100871. 10.1016/j.dcn.2020.100871

Muenssinger, J., Matuz, T., Schleger, F., Kiefer-Schmidt, I., Goelz, R., Wacker-Gussmann, A., Birbaumer, N., & Preissl, H. (2013). Auditory habituation in the fetus and neonate: An fMEG study. Developmental Science, 16(2), 287–295. 10.1111/desc.12025

Musacchia, G., Choudhury, N. A., Ortiz-Mantilla, S., Realpe-Bonilla, T., Roesler, C. P., & Benasich, A. A. (2013). Oscillatory support for rapid frequency change processing in infants. Neuropsychologia, 51(13), 2812–2824. 10.1016/j.neuropsychologia.2013.09.006

Näätänen, R., Paavilainen, P., Rinne, T., & Alho, K. (2007). The mismatch negativity (MMN) in basic research of central auditory processing: A review. Clinical Neurophysiology: Official Journal of the International Federation of Clinical Neurophysiology, 118(12), 2544–2590. 10.1016/j.clinph.2007.04.026

Nguyen, T., Bánki, A., Markova, G., & Hoehl, S. (2020). Chapter 1—Studying parent-child interaction with hyperscanning. In S. Hunnius & M. Meyer (Eds.), Progress in Brain Research (Vol. 254, pp. 1–24). Elsevier. 10.1016/bs.pbr.2020.05.003

Nichols, T. E., & Holmes, A. P. (2002). Nonparametric permutation tests for functional neuroimaging: A primer with examples. Human Brain Mapping, 15(1), 1–25. 10.1002/hbm.1058

Nozaradan, S., Mouraux, A., & Cousineau, M. (2017). Frequency tagging to track the neural processing of contrast in fast, continuous sound sequences. Journal of Neurophysiology, 118(1), 243–253. 10.1152/jn.00971.2016

Oostenveld, R., Fries, P., Maris, E., & Schoffelen, J.-M. (2011). FieldTrip: Open source software for advanced analysis of MEG, EEG, and invasive electrophysiological data. Computational Intelligence and Neuroscience, 2011, 156869. 10.1155/2011/156869

Pfeiffer, C., Andersen, L. M., Lundqvist, D., Hämäläinen, M., Schneiderman, J. F., & Oostenveld, R. (2018). Localizing on-scalp MEG sensors using an array of magnetic dipole coils. PLOS ONE, 13(5), e0191111. 10.1371/journal.pone.0191111

Pfeiffer, C., Ruffieux, S., Andersen, L. M., Kalabukhov, A., Winkler, D., Oostenveld, R., Lundqvist, D., & Schneiderman, J. F. (2020). On-scalp MEG sensor localization using magnetic dipole-like coils: A method for highly accurate co-registration. NeuroImage, 212, 116686. 10.1016/j.neuroimage.2020.116686

QZFM Gen-3 – QuSpin. (n.d.). Retrieved March 25, 2024, from http://quspin.com/products-qzfm/

Regan, D. (1982). Comparison of transient and steady-state methods. Annals of the New York Academy of Sciences, 388, 45–71. 10.1111/j.1749-6632.1982.tb50784.x

Rhodes, N., Sato, J., Safar, K., Amorim, K., Taylor, M. J., & Brookes, M. J. (2024). Paediatric Magnetoencephalography and its Role in Neurodevelopmental Disorders. *The British Journal of Radiology*, tqae123. 10.1093/bjr/tqae123

Rier, L., Rhodes, N., Pakenham, D., Boto, E., Holmes, N., Hill, R. M., Rivero, G. R., Shah, V., Doyle, C., Osborne, J., Bowtell, R., Taylor, M. J., & Brookes, M. J. (2024). The neurodevelopmental trajectory of beta band oscillations: An OPM-MEG study. eLife, 13. 10.7554/eLife.94561.1

Seymour, R. A., Alexander, N., Mellor, S., O’Neill, G. C., Tierney, T. M., Barnes, G. R., & Maguire, E. A. (2021). Using OPMs to measure neural activity in standing, mobile participants. NeuroImage, 244, 118604. 10.1016/j.neuroimage.2021.118604

Seymour, R. A., Alexander, N., Mellor, S., O’Neill, G. C., Tierney, T. M., Barnes, G. R., & Maguire, E. A. (2022). Interference suppression techniques for OPM-based MEG: Opportunities and challenges. NeuroImage, 247, 118834. 10.1016/j.neuroimage.2021.118834

Shibasaki, H., & Miyazaki, M. (1992). Event-related potential studies in infants and children. Journal of Clinical Neurophysiology: Official Publication of the American Electroencephalographic Society, 9(3), 408–418. 10.1097/00004691-199207010-00007

Tierney, T. M., Holmes, N., Mellor, S., López, J. D., Roberts, G., Hill, R. M., Boto, E., Leggett, J., Shah, V., Brookes, M. J., Bowtell, R., & Barnes, G. R. (2019). Optically pumped magnetometers: From quantum origins to multi-channel magnetoencephalography. NeuroImage, 199, 598–608. 10.1016/j.neuroimage.2019.05.063

Vigário, R., Särelä, J., Jousmäki, V., Hämäläinen, M., & Oja, E. (2000). Independent component approach to the analysis of EEG and MEG recordings. IEEE Transactions on Bio-Medical Engineering, 47(5), 589–593. 10.1109/10.841330

Wang, Q., Zhu, G.-P., Yi, L., Cui, X.-X., Wang, H., Wei, R.-Y., & Hu, B.-L. (2019). A Review of Functional Near-Infrared Spectroscopy Studies of Motor and Cognitive Function in Preterm Infants. Neuroscience Bulletin, 36(3), 321–329. 10.1007/s12264-019-00441-1

Wens, V. (2023). Exploring the limits of MEG spatial resolution with multipolar expansions. NeuroImage, 270, 119953. 10.1016/j.neuroimage.2023.119953

Wunderlich, J. L., & Cone-Wesson, B. K. (2006). Maturation of CAEP in infants and children: A review. Hearing Research, 212(1), 212–223. 10.1016/j.heares.2005.11.008

Wunderlich, J. L., Cone-Wesson, B. K., & Shepherd, R. (2006). Maturation of the cortical auditory evoked potential in infants and young children. Hearing Research, 212(1–2), 185–202. 10.1016/j.heares.2005.11.010

Zahran, S., Mahmoudzadeh, M., Wallois, F., Betrouni, N., Derambure, P., Le Prado, M., Palacios-Laloy, A., & Labyt, E. (2022). Performance Analysis of Optically Pumped 4He Magnetometers vs. Conventional SQUIDs: From Adult to Infant Head Models. Sensors, 22(8), Article 8. 10.3390/s22083093

Zetter, R., Iivanainen, J., & Parkkonen, L. (2019). Optical Co-registration of MRI and On-scalp MEG. Scientific Reports, 9(1), 5490. 10.1038/s41598-019-41763-4

