## Supplemental Material for "Pushing the boundaries of MEG based on optically pumped magnetometers towards early human life"

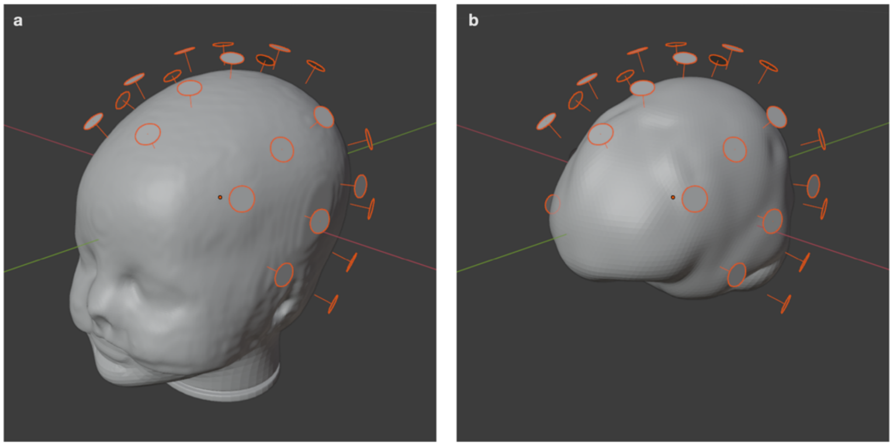


**Figure S1. Co-registration procedure**. Position and radial axis orientation of the sensors with respect (**a**) to the 3D-printed head model, which comes from the infant MNI MRI template and (**b**) to the inner skull surface used for the source reconstruction analysis, which is the adult MNI MRI inner skull surface, shrunk to contain the template BCP brain as snugly as possible.

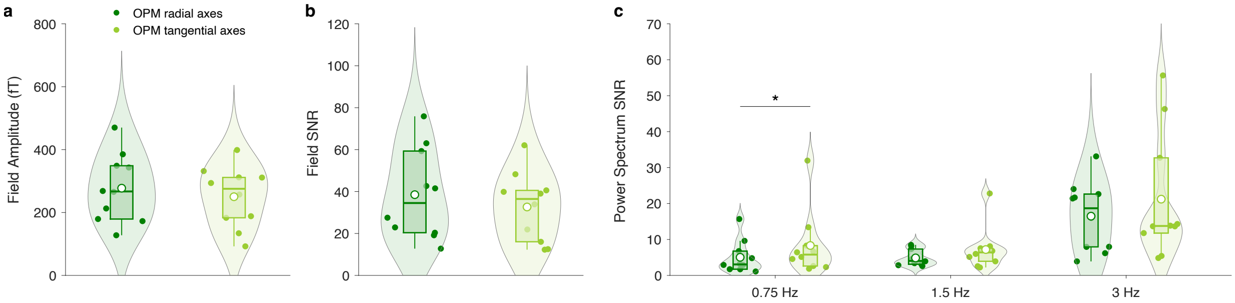


**Figure S2. Comparison of the responses recorded by the OPMs on the radial and tangential axes for the 10 participants for whom triaxial sensors were used**. (**a**) Amplitude of the evoked responses, (**b**) SNR of the evoked responses and (**c**) power spectrum SNR at the frequencies of interest. The white circles represent the mean of the data. The statistical significance of the comparisons between the axial and tangential axes is indicated above the violin plots. *$p < 0.05$.

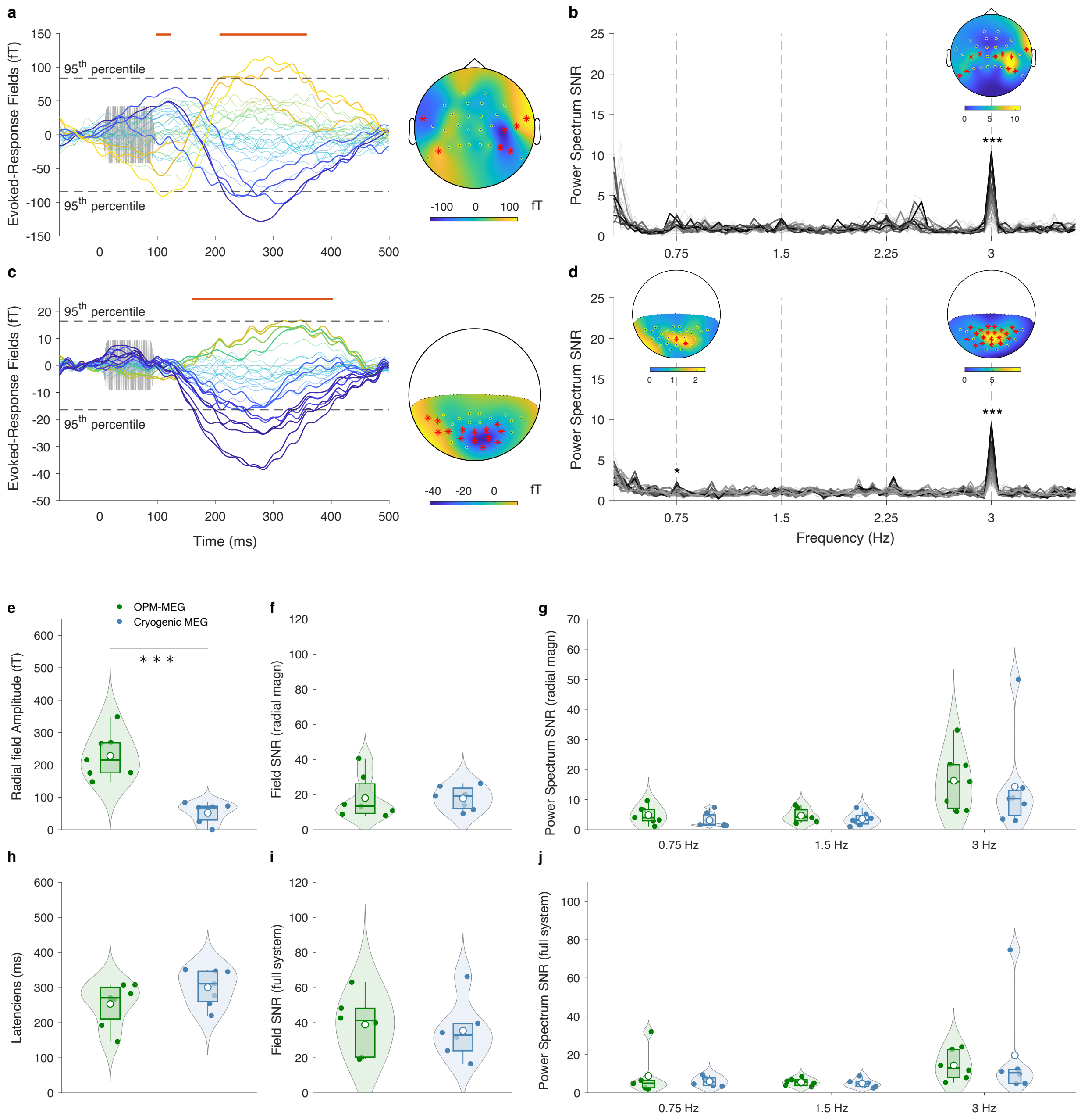


**Figure S3. Comparison of the OPM-MEG and cryogenic MEG system**. **Left.** Radial magnetic fields evoked in response to single tones measured with (**a**) OPM-MEG and (**c**) cryogenic MEG. The plots are the same as in Fig. 2. Unsurprisingly, given that the infants were lying on their sides in the cryogenic MEG setup, we observed a single dipole in the posterior cryogenic sensors. We also show a comparison of (**e**) the amplitudes of the evoked response of the radial magnetometers, (**f**) their SNRs and (**h**) their latencies. (**i**) For completeness we also show a comparison of the field SNR when considering all channels of both systems (i.e., including tangential axes for the OPM-MEG and gradiometers for cryogenic MEG). **Right.** Radial magnetic field spectrum SNR for the steady-state response obtained with (**b**) OPM-MEG and (**d**) cryogenic MEG. All is the same as in Fig. 3. The power spectrum SNRs are compared at 0.75 Hz and 3 Hz for the radial magnetometers (**g**) and for the full systems (**j**). For the radial magnetometers, the comparisons were done for the 7 participants who underwent the experiments with both modalities (paired tests), while for the full system the comparisons were done for 6 subjects that had tri-axial OPM recordings and 6 other subjects that had cryogenic MEG recordings (unpaired tests). The white circles represent the mean of the data. The statistical significance of the comparisons between OPM and cryogenic MEG is indicated above the violin plots. ***$p < 0.001$.

| | Evoked response paradigm | | Oddball paradigm | |
|---|---|---|---|---|
| | OPM-MEG | Cryogenic MEG | OPM-MEG | Cryogenic MEG |
| Length of recorded data (s) | 575 ± 227 | 544 ± 143 | 570 ± 244 | 468 ± 171 |
| Length of artefact-free data (s) | 450 ± 213 | 492 ± 151 | 458 ± 235 | 428 ± 165 |
| Number of periods with artifacts | 11 ± 6 | 5 ± 4 | 11 ± 3 | 4 ± 4 |
| Mean length of periods with artifacts (s) | 10 ± 3 | 9 ± 1 | 9 ± 3 | 7 ± 2 |
| Length of artifact-free data after z-score rejection (s) | 404 ± 191 | 484 ± 152 | N/A | N/A |

**Table S1. Duration of recorded data and periods with artifact.** For both paradigms and both MEG systems, the mean and standard deviation of the total length of the recordings, the total length of the artefact-free parts, the number of periods with artifacts, and their averaged lengths are presented. For the evoked response paradigm, we also show the amount of data remaining after applying the z-score rejection as described in the Methods section.

| | Number of trials for the Evoked response paradigm | | | | Number of 20-s epochs for the Oddball paradigm | | | |
|---|---|---|---|---|---|---|---|---|
| | OPM-MEG | | Cryogenic MEG | | OPM-MEG | | Cryogenic MEG | |
| | Recorded | Accepted | Recorded | Accepted | Recorded | Accepted | Recorded | Accepted |
| Mean | 1038 | 694 | 954 | 849 | 28 | 16 | 23 | 19 |
| SD | 412 | 328 | 271 | 295 | 12 | 11 | 8 | 7 |
| Min | 547 | 230 | 650 | 501 | 15 | 3 | 18 | 13 |
| Max | 1751 | 1223 | 1335 | 1325 | 54 | 36 | 43 | 35 |

**Table S2. Number of trials (for the Evoked response paradigm) and 20-s epochs (for the Oddball paradigm).** Both the number of trials recorded and accepted (mean, standard deviation, minimum and maximum) are presented for the two different MEG systems.

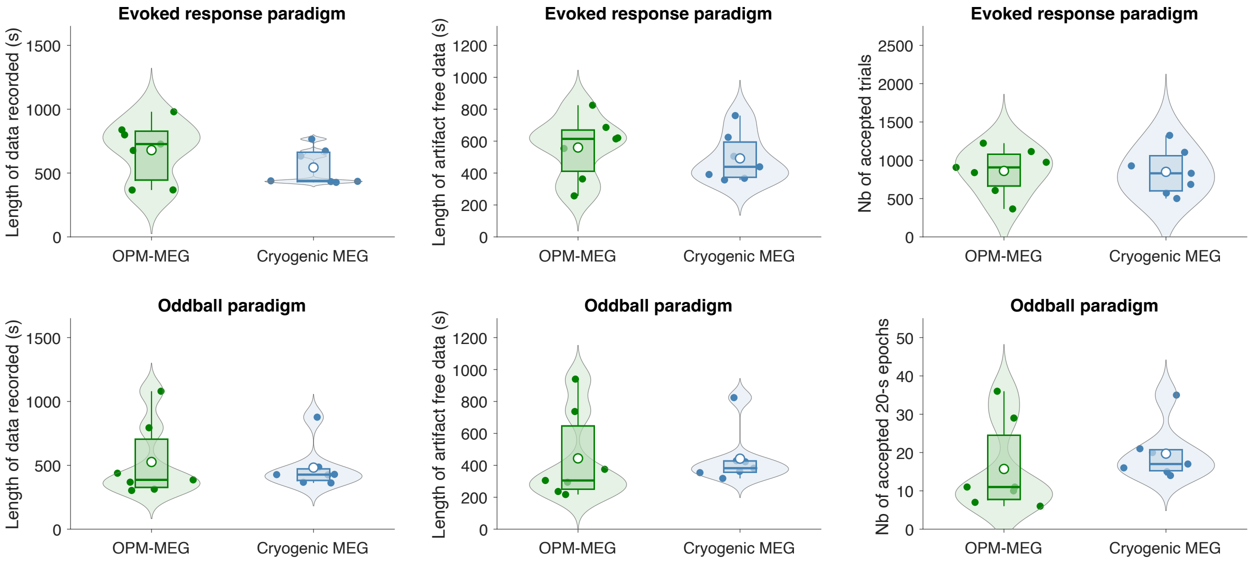


**Figure S4. Comparison of the amount of data between the OPM-MEG and cryogenic systems**. The comparisons for the total amount of recorded data, the total amount of artifact free data, the number of accepted trials (evoked response paradigm) and the number of accepted 20-s epochs (Oddball paradigm) are shown. None of the differences is statistically significant.
